## Supplemental figures 1-7 and legends for "Ubiquitin Ligase ITCH Regulates Life Cycle of SARS-CoV-2 Virus"

**Expanded View Figure Legends**

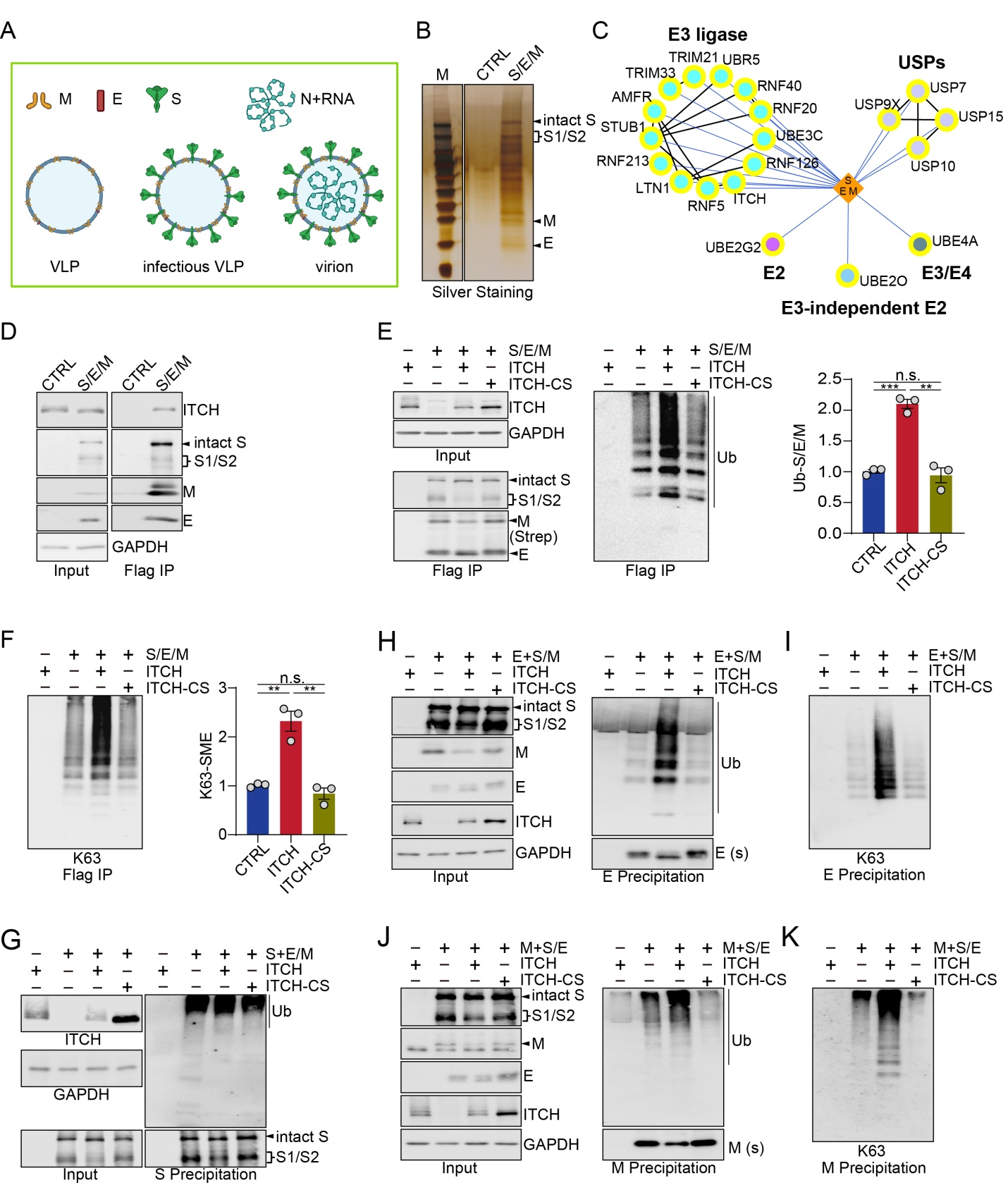

**Figure supplement 1. ITCH promotes ubiquitination of SARS-CoV-2 structural proteins.**

**(A)**	Components of SARS-CoV-2 VLP, infectious VLP, and infectious virion.

**(B)**	Lysates of HEK293 cells expressing Flag-2×Strep-tagged S/E/M proteins were subjected to Flag immunoprecipitation (IP) and Strep affinity precipitation (AP), followed by detection of proteins by silver staining.

**(C)**	The interaction network between structural proteins and ubiquitination related proteins as identified by the proteomic analysis.

**(D)**	Lysates from HEK293 cells expressing Flag-2×Strep tagged S/E/M were subjected to Flag IP. Immunoblotting analysis showed that ITCH was pulled down by the structural proteins.

**(E**, **F)** Denatured lysates from HEK293 cells expressing HA-tagged ubiquitin (Ub), Flag-tagged S, Flag-2×Strep-tagged E/M with ITCH or ITCH-CS were subjected to Flag IP, followed by immunoblotting against ubiquitin (Ub), Ub-K63, and the structural proteins (S antibody for S and Strep antibody for E/M). The immunoblotting analysis indicated that ITCH, but not the inactive ITCH-CS, promoted the ubiquitination of one or more components of the complex formed by the S/E/M structural proteins, through K63 polyubiquitin chains (n=3).

**(G)**	Denatured lysates from HEK293 cells expressing HA-tagged ubiquitin (Ub), s-tag-fused E/M, and Flag-tagged S with ITCH or ITCH-CS were subjected to Flag IP. Immunoblotting analysis of the immunoprecipitates with ubiquitin antibody showed that the ubiquitination of S protein was not altered by ITCH.

**(H**, **I)** Denatured lysates from HEK293 cells expressing HA-tagged ubiquitin (Ub), Flag-tagged S, Flag-2×Strep-tagged M, and s-tag-fused E with ITCH or ITCH-CS were subjected to s-tag AP. Immunoblotting analysis of the precipitates with ubiquitin (Ub) or Ub-K63 antibodies showed that, in the presence of M/S, the ubiquitination of E was increased by ITCH, but not the inactive ITCH-CS, through K63-linked chains.

**(J**, **K)** Denatured lysates from HEK293 cells expressing HA-tagged ubiquitin (Ub), Flag-tagged S, Flag-2×Strep-tagged E, and s-tag-fused M with ITCH or ITCH-CS were subjected to s-tag AP. Immunoblotting analysis of the precipitates with ubiquitin (Ub) or Ub-K63 antibodies showed that, in the presence of E/S, the ubiquitination of M was increased by ITCH, but not the inactive ITCH-CS, through K63-linked chains.

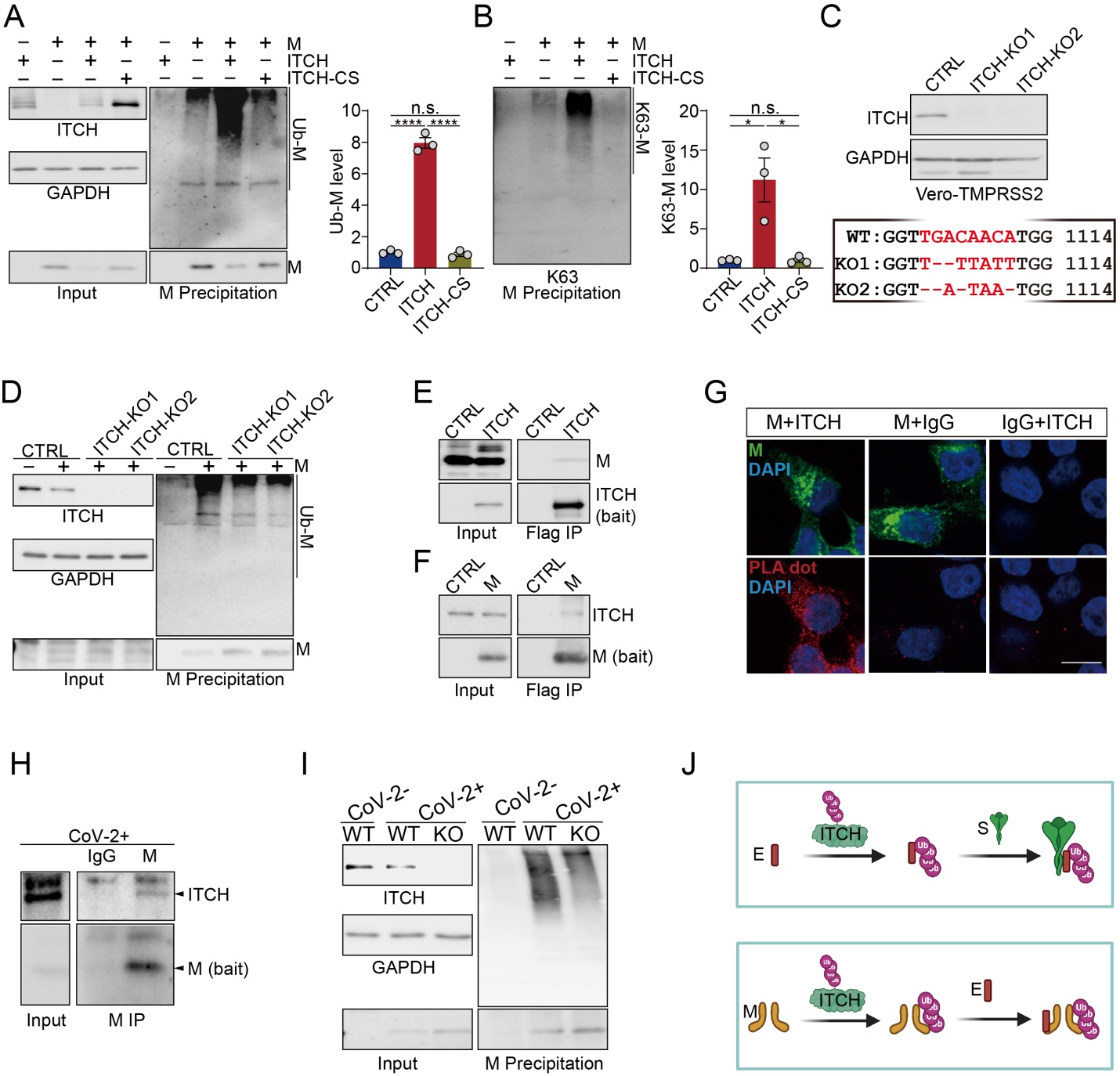

**Figure supplement 2. ITCH interacts with M and promotes its ubiquitination.**

**(A**, **B)** Denatured lysates from HEK293 cells expressing HA-tagged ubiquitin (Ub), Flag-tagged M with ITCH or ITCH-CS were subjected to Flag IP. Immunoblotting analysis of the immunoprecipitates with ubiquitin (Ub) or Ub-K63 antibodies showed that the ubiquitination of M was increased by ITCH, but not the inactive ITCH-CS, through K63-linked chains WT(n=3).

**(C)**	Validation of the vT2-ITCH-KO cell lines through immunoblotting of ITCH and sequencing of the sgRNA-targeted DNA region.

**(D)**	Denaturing Strep AP using lysates from vT2-WT and vT2-ITCH-KO cells expressing HA tagged ubiquitin (Ub), Flag-2×Strep tagged M was performed. A decrease in the ubiquitination of M was shown in the ITCH-KO cells.

**(E)**	Non-denatured lysates from HEK293 cells expressing s-tag-fused M with Flag-tagged ITCH were subjected to Flag IP, showing the specific pull-down of M by ITCH.

**(F)**	Non-denatured lysates from HEK293 cells expressing RFP or Flag-tagged M were subjected to Flag IP, showing the specific pull-down of endogenous ITCH by M protein.

**(G)** The proximal colocalization of ITCH with M was analyzed in HEK293 cells by PLA using a rabbit anti-Flag antibody paired with a mouse anti-ITCH antibody. Negative controls included non-specific mouse IgG or rabbit IgG paired with the Flag or ITCH antibody, respectively (n>15 cells). Scale bar, 10 μm.

**(H)**	Lysates from vT2 cells infected with SARS-CoV-2 at 1 MOI for 10 h were subjected to M IP, followed by immunodetection using antibody against ITCH. An ITCH signal was clearly observed in the M immunoprecipitates.

**(I)**	M IP was carried out using denatured lysates from vT2-WT and vT2-ITCH-KO cells infected with SARS-CoV-2 at 1 MOI for 10 hours, Immunoblotting analysis revealed a reduction in ubiquitinated M in the absence of ITCH.

**(J)** Models depicting the role of ITCH in promoting mutual interactions among the SARS-CoV-2 structural proteins.

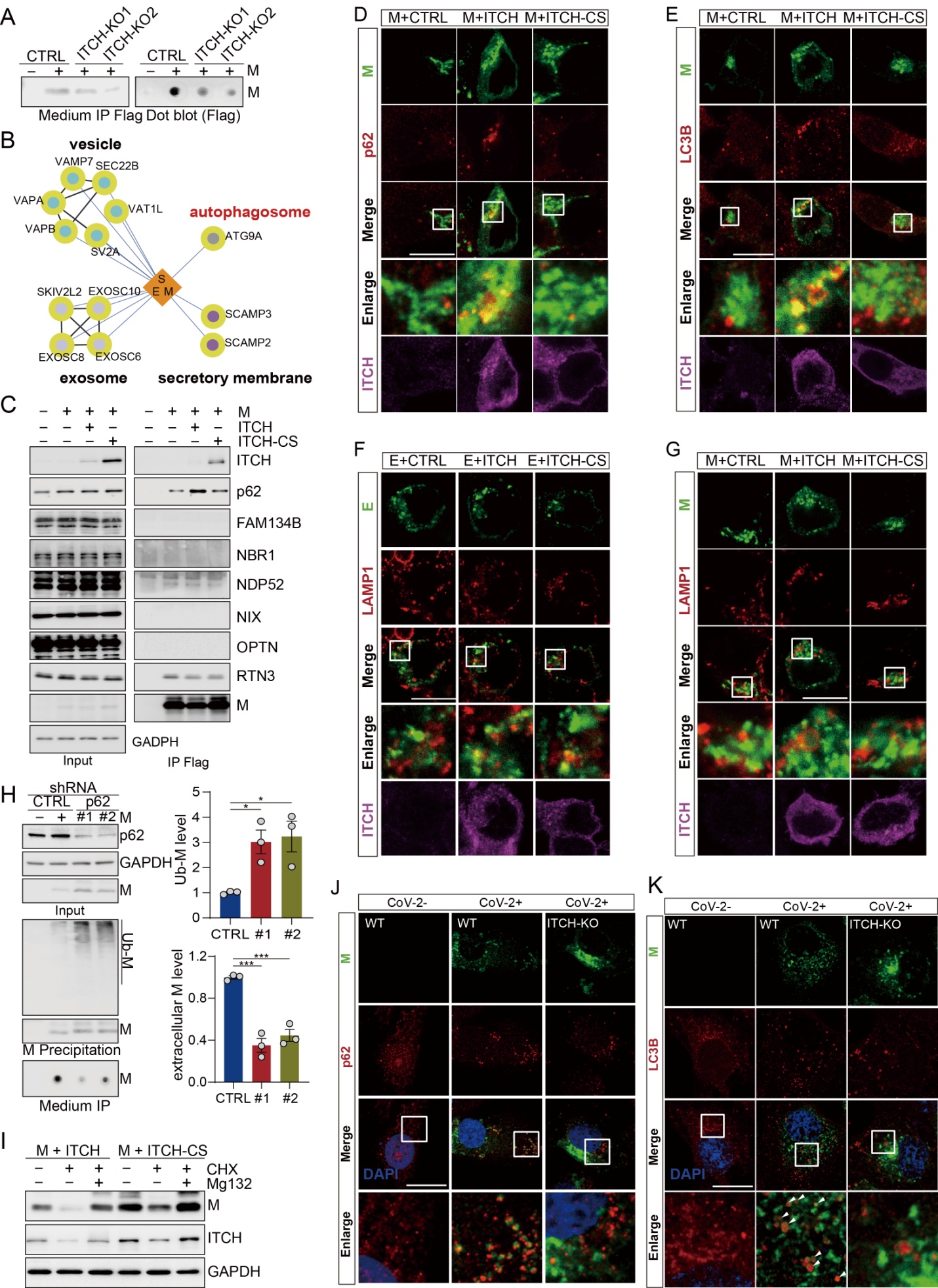

**Figure supplement 3. ITCH promotes the trafficking of M from TGN to p62-dependent autophagosomes and enhances its secretion.**

**(A)**	Culture media from vT2-WT and vT2-ITCH-KO cells expressing HA-tagged ubiquitin (Ub) and Flag-2×Strep tagged M were harvested for the Strep AP. Proteins were then visualized by immunoblot or dot blot analysis, showing a decrease of the extracellular M protein in the absence of ITCH.

**(B)**	The interaction network between the SARS-CoV-2 structural proteins and secretion host proteins as identified by the proteomic analysis.

**(C)**	Co-IP analysis of lysates from HEK293 cells expressing Flag-tagged M with ITCH or ITCH-CS showed that ITCH, but not the inactive ITCH-CS, promoted the specific interaction between M and p62.

**(D-E)** Immunofluorescence analysis of HEK293 cells expressing Flag-tagged M with ITCH or ITCH-CS showed that ITCH, but not the inactive ITCH-CS, enhanced the colocalization between M and p62 (D) or LC3B (E), while the colocalization between M and LAMP1 was not altered (E). Scale bar, 10 μm.

**(F, G)** HEK293 cells were transfected with Flag-tagged E or M with ITCH or ITCH-CS. 24 h later, cells were analyzed by immunofluorescence with Flag, ITCH and LAMP1 antibodies. ITCH expression did not alter the colocalization of LAMP1 with E (F) or M (G). Scale bar, 10 μm.

**(H)**	Flag IP was conducted to precipitate M protein from culture media and denatured lysates of control (CTRL) and p62 knockdown (#1, #2) from HEK293 cells expressing HA-tagged ubiquitin (Ub) and Flag-tagged M, showing that the levels of both unmodified and ubiquitinated intracellular M were increased by p62 depletion, with a concurrent decrease in the level of extracellular M (n=3).

**(I)**	HEK293 cells expressing M together with ITCH or ITCH-CS were cultured for 12 h, followed by treatment with CHX (10 μg/ml) and MG132 (2 nM) or DMSO for 24 h. Immunoblotting revealed that proteasome inhibition prevented M turnover but failed to restore M levels reduced by ITCH expression.

**(J, K)** vT2-WT and vT2-ITCH-KO cells infected with SARS-CoV-2 at 1 MOI for 10 h were subjected to immunofluorescence with M and p62 or LC3B antibodies. Colocalization between E and p62 (H) or LC3B (I) was significantly reduced with ITCH ablation. Scale bar, 10 μm.

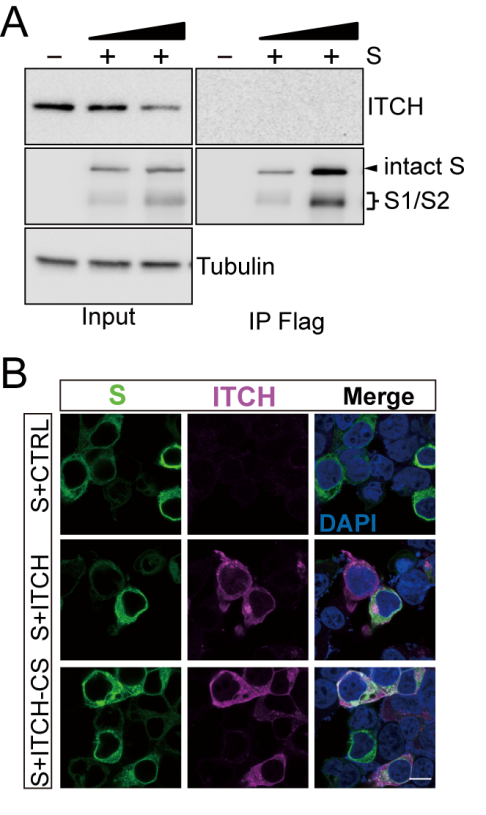

**Figure supplement 4. ITCH does not associate with S and does not affect its subcellular localization.**

**(A)**	Non-denatured lysates from HEK293 cells expressing Flag-tagged S protein were subjected to Flag IP, showing that ITCH signal was not detected in the S immunoprecipitates.

**(B)**	Subcellular localization analysis of Flag-tagged S in HEK293 cells expressing ITCH or ITCH-CS showed that ITCH did not regulate the subcellular localization of S protein. Scale bar, 10 μm.

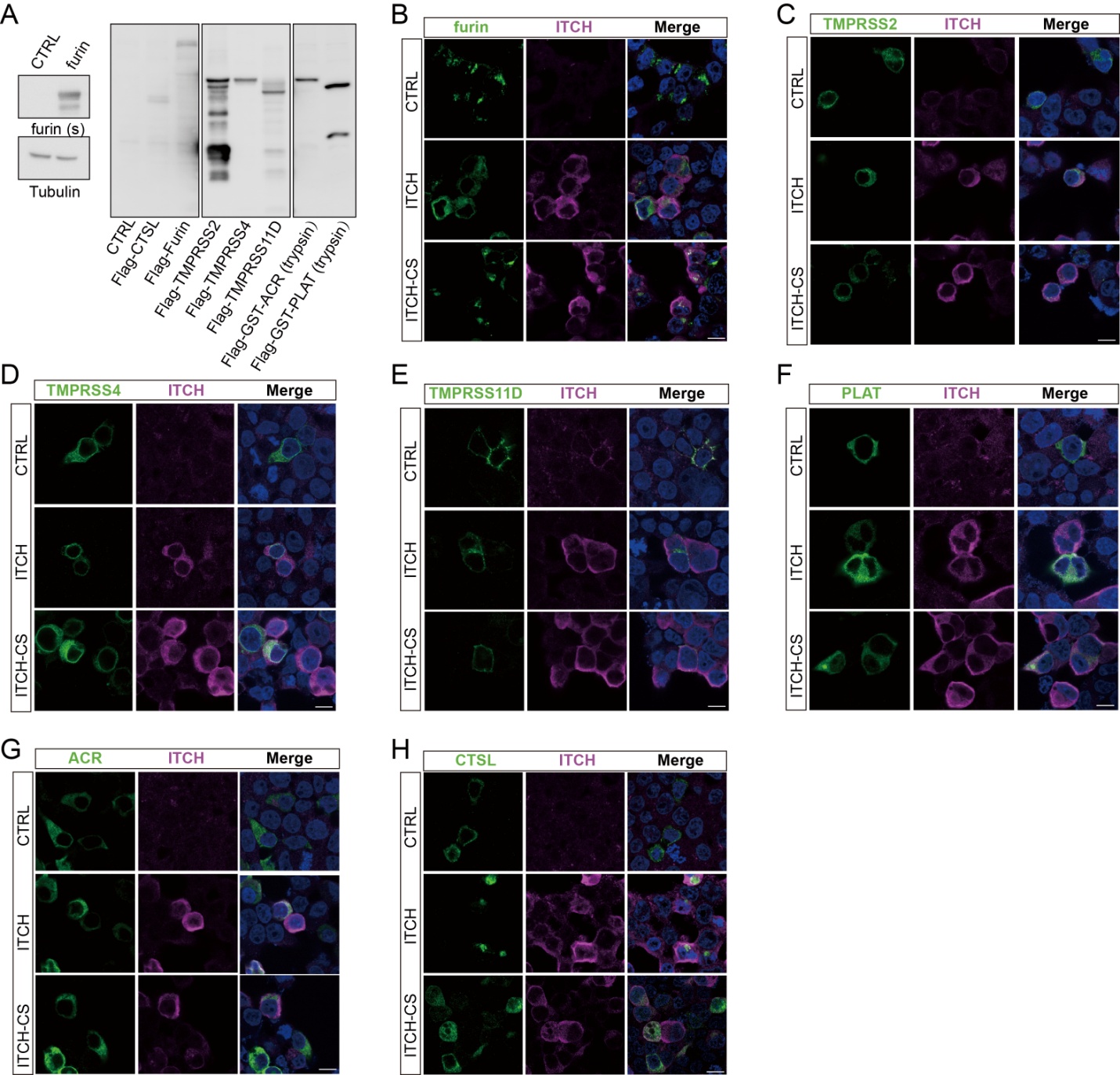

**Figure supplement 5. ITCH does not affect subcellular localization of S-related TMPRSS and trypsin proteases.**

**(A)**	Validation of the expression of various TMPRSS and trypsin proteases in HEK293 cells.

**(B-H)** HEK293 cells were transfected with the indicated Flag-tagged proteases together with ITCH or ITCH-CS. Immunofluorescence analysis revealed no detectable changes in the subcellular distribution of these proteases upon ITCH overexpression, with the exception of furin and CTSL. Scale bar, 10 μm.

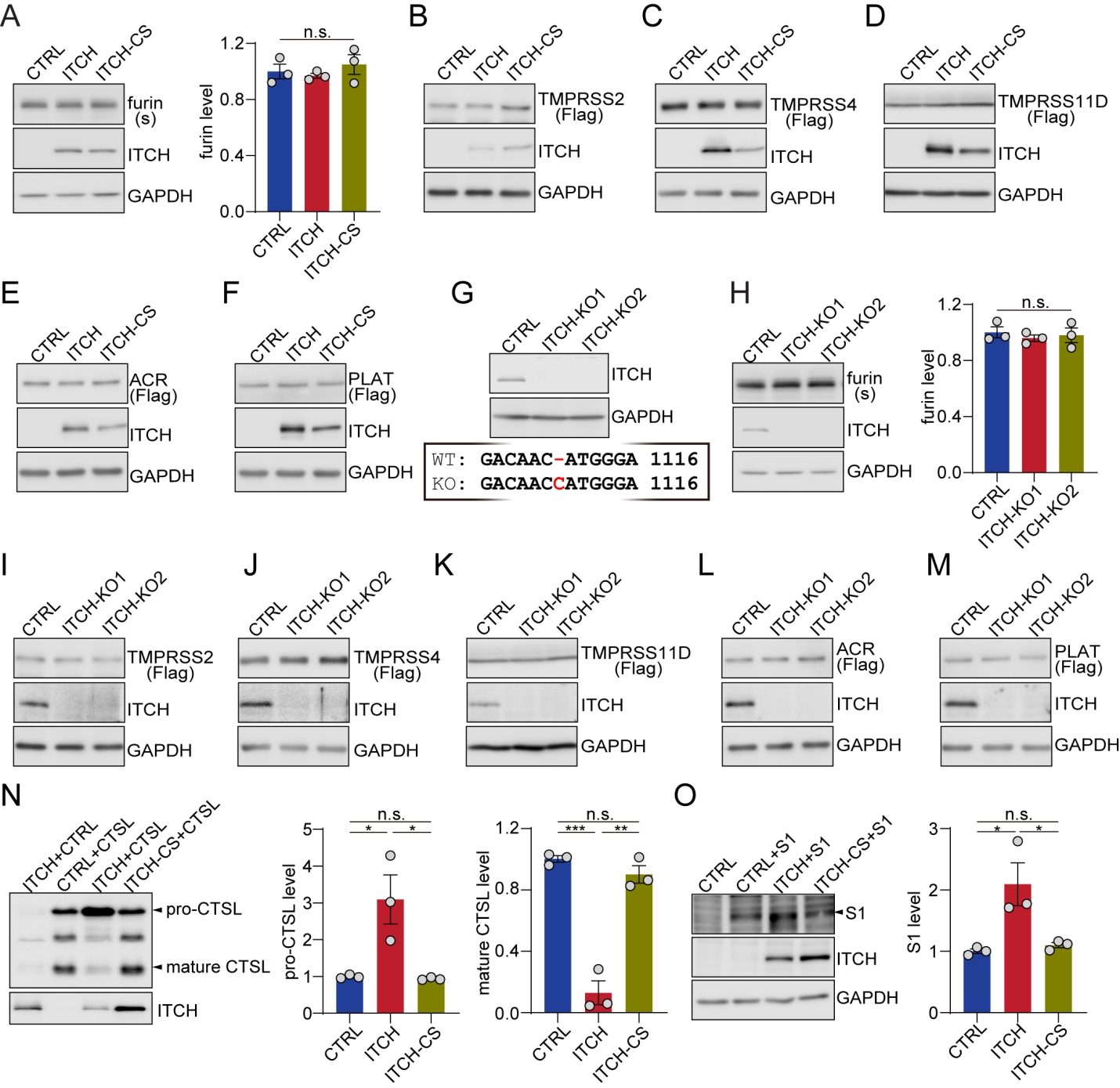

**Figure supplement 6. ITCH does not affect the levels of TMPRSS and trypsin proteases.**

**(A)**	The levels of furin in HEK293 cells expressing WT or inactivated ITCH were analyzed by immunoblotting, showing that the level of furin was not altered by ITCH overexpression (n=3).

**(B-F)** HEK293 cells were transfected with Flag-tagged TMPRSS proteases (TMPRSS2, TMPRSS4, TMPRSS11D) and trypsins (ACR, PLAT) with ITCH or ITCH-CS. No change in the levels of these proteases was observed upon ITCH overexpression.

**(G)**	Validation of the HEK293 ITCH-KO cell lines through immunoblotting of ITCH and sequencing of the sgRNA-targeted DNA region.

**(H)**	The levels of furin in HEK293 WT and ITCH-KO cells were analyzed by immunoblotting, showing that the level of furin was not altered by ITCH ablation (n=3).

**(I-M)** HEK293 WT and ITCH-KO cells were transfected with Flag-tagged TMPRSS and trypsin proteases, followed by immunoblotting with Flag or ITCH antibodies, showing that ITCH ablation did not change the levels of these proteases.

**(N)**	Immunoblotting analysis of lysates from HEK293 cells expressing CTSL and ITCH or ITCH-CS showed that ITCH, but not ITCH-CS, decreased the level of mature CTSL with a concurrent increase of pro-CTSL (n = 3).

**(O)**	Immunoblotting analysis of the S1 subunit using lysates from HEK293 cells expressing N-terminally Flag-tagged S1 and ITCH or ITCH-CS showed that ITCH, but not ITCH-CS, increased the level of S1 (n = 3).

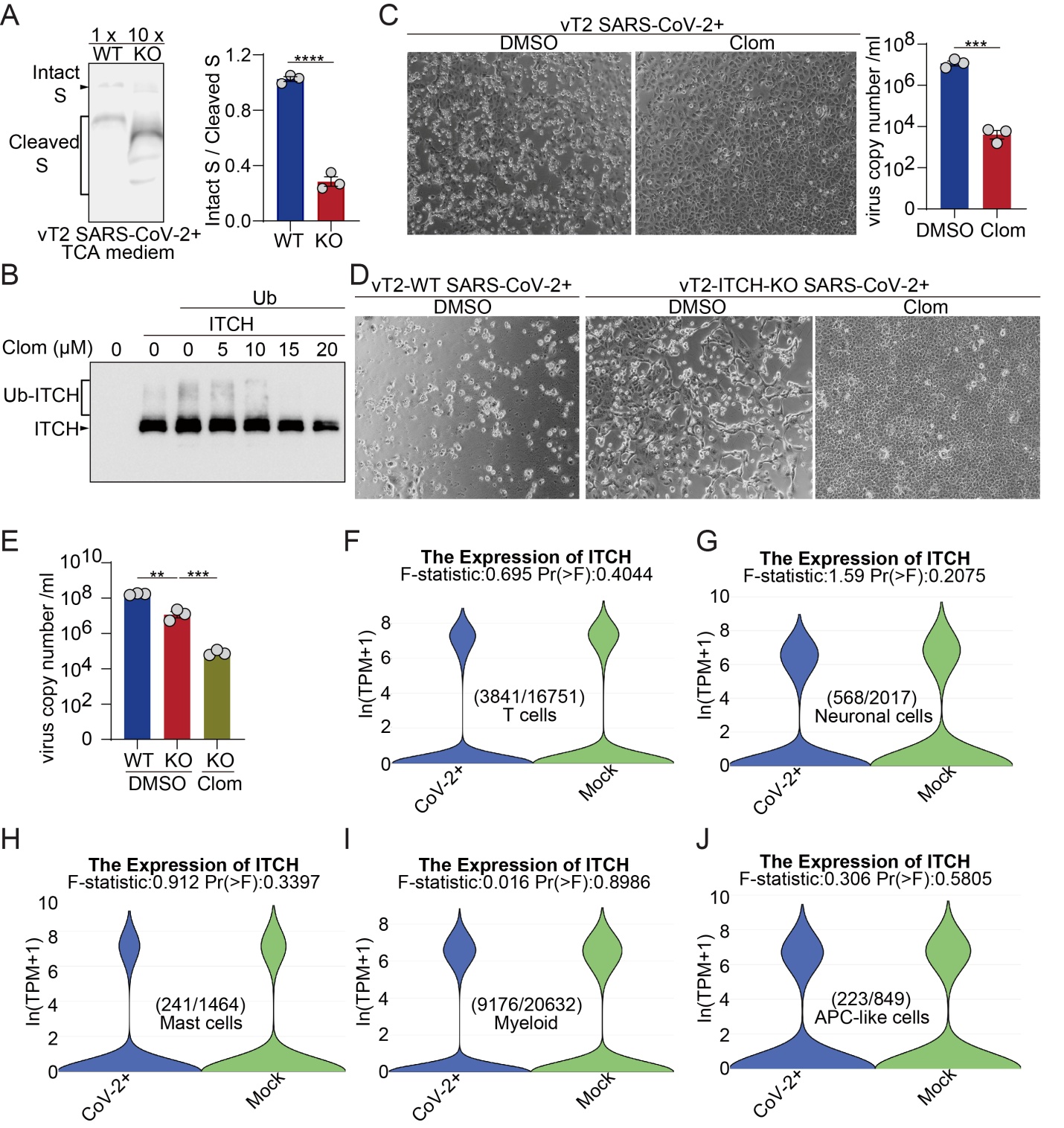

**Figure supplement 7.** **Pharmacological inhibition of ITCH suppresses SARS-CoV-2 production.**

**(A)**	Culture media from vT2-WT and vT2-ITCH-KO cells infected with SARS-CoV-2 for 48 h were collected and subjected to TCA precipitation. S proteins (1× input from WT and 10× input from ITCH-KO samples) were analyzed by immunoblotting, revealing a marked increase in S protein cleavage in the absence of ITCH.

**(B)**	Immunoblotting analysis of vT2 cells expressing ubiquitin and Flag-tagged ITCH and treated with various concentrations of Clom (18 h) showed that Clom inhibited the activity of ITCH, as indicated by its self-ubiquitination, at the concentration of 15 μM or above.

**(C)**	Analysis of vT2 cells infected with SARS-CoV-2 at 0.0001 MOI showed that the SARS-CoV-2-induced CPE and virus production as indicated by virus copy number (48 hpi) were significantly inhibited by the Clom treatment (n=3).

**(D, E)**	Analysis of vT2-WT and vT2-ITCH-KO cells infected with SARS-CoV-2 at 0.0001 MOI showed that SARS-CoV-2-induced viral copy numbers in the culture medium (48 hpi) were reduced by several hundred-fold in vT2-ITCH-KO cells treated with Clom (n=3).

**(F-J)** ITCH expression analysis using published single-nucleus RNA sequencing data from the COVID-19 patients and control individuals revealed no significant difference in ITCH mRNA levels in T cells, neurons, mast cells, myeloid cells, or APC-like cells in the lungs with or without SARS-CoV-2 infection.
